# Computable Genomes: A Layered Computational Framework for Non-Coding DNA and Cellular Behavior

**DOI:** 10.64898/2026.11.10.765552

**Authors:** Alejandra Martin Sevilla, Lurong Pan

## Abstract

Genome-wide association studies have catalogued tens of thousands of trait-associated loci, most of them in non-coding DNA, yet for most loci the path from nucleotide to phenotype remains correlational. We propose a layered computational decomposition of the genome as a complement to data-driven approaches: sequence as formal language, sequence as information-bearing signal, mobile elements as runtime, compilation to interaction networks through thermodynamic and rule-based models, and execution as cellular and multicellular dynamics. Each layer is defined by its inputs, outputs and formalisms, and we set out where the frame breaks down, including the looseness of algorithmic-information estimates, the triviality of universality results and the failure of string grammars at the chromatin level. Synthetic-biology platforms serve as tests of whether the layers compose, and genomic foundation models and the AlphaGenome Atlas, which predicts effects for all nine billion possible single-nucleotide variants, serve as oracles whose coverage and limits we map onto the framework. We state five falsifiable predictions and test the first, that sequence compressibility carries information about regulatory function beyond composition and conservation, on 38,556 single-nucleotide variants from a saturation-mutagenesis reporter assay of 21 regulatory elements. In a pooled model, local compressibility is associated with measured effect size (standardized β = −0.030, p = 0.0014). The association roughly halves under full control for base composition, loses significance when composition, motif and noise adjustments are combined under locus-level inference (wild cluster bootstrap p = 0.11), and averages zero within loci (random-effects β = −0.0003, p = 0.90). The prediction is therefore not supported at the variant level in these data, which bound the effect a genome-wide test should expect. The argument is not that the genome “is” a program, but that treating it as one supplies formal tools whose predictions can be checked.

## 1. Introduction

Genomics has accumulated more sequence than mechanism. Genome-wide association studies have catalogued tens of thousands of trait-associated loci, the great majority lying in non-coding DNA [1], yet for most of these loci the path from nucleotide to phenotype remains correlational rather than causal. The ENCODE registry of candidate cis-regulatory elements (cCREs) grew from 926,535 human and 339,815 mouse elements in Phase III, covering approximately 7.9% and 3.4% of the respective genomes [2], to about 2.37 million human and more than 900,000 mouse elements in the ENCODE 4 Registry [66]. The ENCODE 4 Functional Characterization Centers assembled the largest collective dataset of non-coding CRISPR screens to date, jointly analyzing over 100 perturbation screens to standardize cCRE validation methodology [3]. These numbers confirm that the non-coding genome is not junk but a vast, structured regulatory architecture whose logic remains largely undeciphered.

The standard response, train a larger model on more data, has produced impressive predictions but limited mechanistic insight [4,67]. The most dramatic example is Evo 2, a 40-billion-parameter genomic foundation model developed by the Arc Institute and NVIDIA with academic collaborators, trained on 9.3 trillion nucleotides spanning all domains of life using a StripedHyena 2 backbone with a 1-megabase context length [5]. Evo 2 recovers known regulatory grammars (CRISPR arrays, tRNA structures, transcription-factor binding motifs) from raw sequence without explicit annotation, and Goodfire’s mechanistic-interpretability collaboration extracted sparse-autoencoder features for alpha-helices, beta-sheets, and tRNAs directly from Evo 2’s activations [6]. These results demonstrate that structured regulatory codes are learnable from sequence alone, but they do not explain why those codes work or how they compose.

This paper takes the complementary route. It asks what we can recover about the function of non-coding DNA by treating the genome as the input to a computation, using formalisms that were designed to reason about computation in the first place: formal language theory [7,8], algorithmic information theory [9], statistical mechanics [10], and automata theory [11]. The thesis is narrower than the slogan “the genome is a program,” which we regard as a useful framing device but not a defensible mechanistic claim [68,69]. We argue instead that the genome admits a layered computational decomposition, sequence, information content, mobile-element runtime, compilation to interaction networks, and execution as cellular and multicellular dynamics, and that the productive work happens at the interfaces between layers (Figure 1).

**Figure 1.**
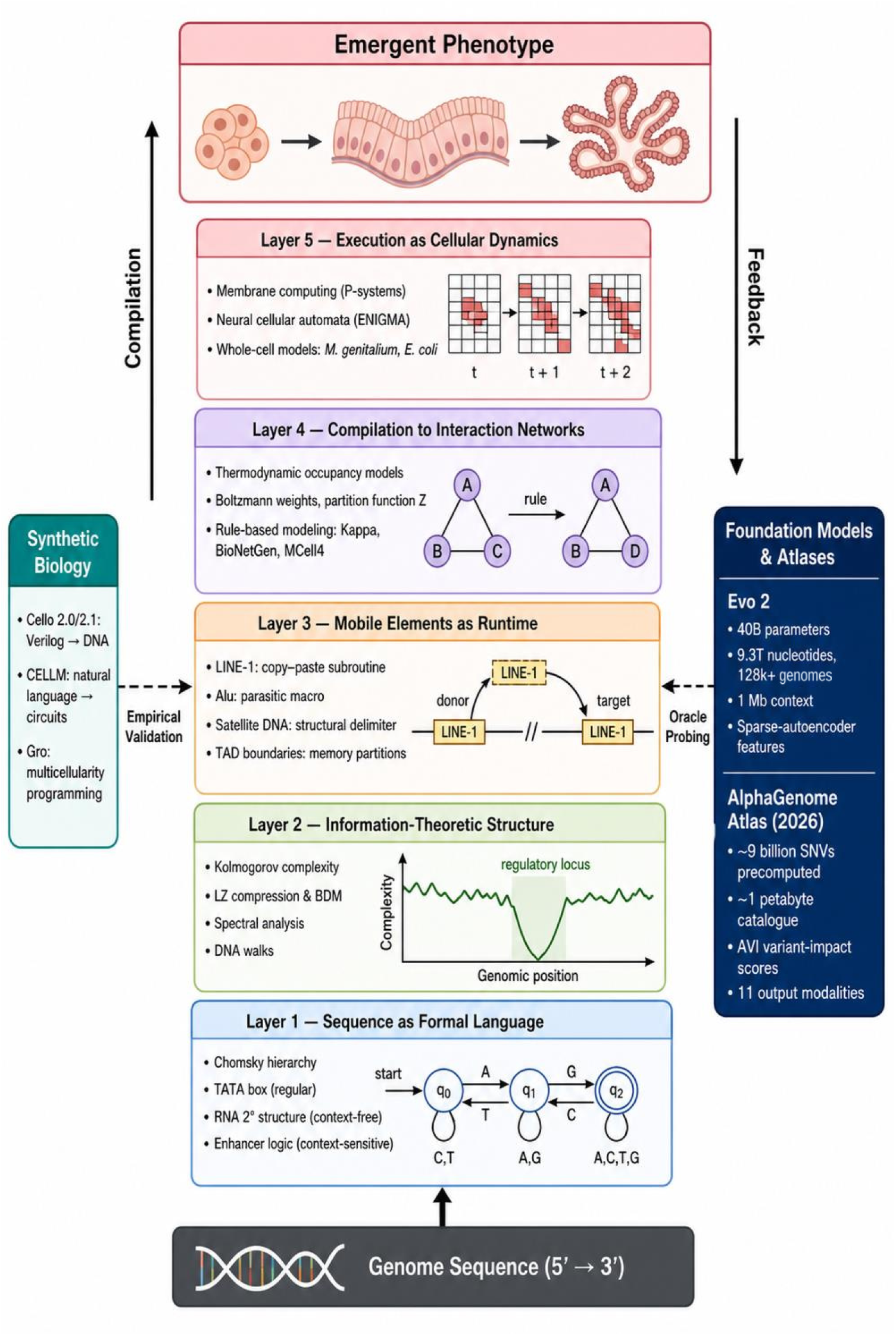
A layered computational decomposition of the genome. Genomic sequence (bottom) is parsed as a formal language (Layer 1), quantified information-theoretically (Layer 2), historically rewritten by mobile elements acting as a runtime (Layer 3), compiled into interaction networks by thermodynamic occupancy and rule-based models (Layer 4), and executed as cellular and multicellular dynamics (Layer 5), yielding emergent phenotype (top). Synthetic-biology platforms (left: Cello 2.0/2.1, CELLM, Gro) supply empirical validation of the stack, while genomic foundation models (Evo 2) and the AlphaGenome Atlas precomputed variant-effect catalogue (right) serve as oracles for probing it. Figure created with Biomni (Phylo).

To keep the decomposition testable, we use the computational vocabulary only as shorthand for maps between measurable objects. Layer 1 maps a sequence to the grammar classes needed to describe its recognized features; layer 2 maps it to information-theoretic descriptors such as compressibility, spectra and walk statistics; layer 3 maps the history of insertions to the regulatory sequence present in a given genome; layer 4 maps sequence and factor concentrations to binding-state probabilities and reaction rules; and layer 5 maps those rules to cellular and tissue trajectories. We say that the stack composes for a perturbation when chaining the layer-specific models predicts its measured downstream effect at least as well as a model fitted directly to that effect, and that it fails to compose when it does not. Terms such as runtime, macro or compilation label these maps; they are not mechanistic claims.

We state five predictions, each with a falsification condition, in Section 11, and test the first on measured data in Section 9: (1) sequence compressibility carries information about regulatory function beyond composition and conservation; (2) SCFG-derived structural priors improve non-coding RNA variant classification; (3) thermodynamic occupancy models extend predictively to simple eukaryotic systems given measured parameters; (4) rule-based reconstruction identifies the causal chain from variant to phenotype; and (5) small gene-regulated NCAs reproduce developmental robustness.

### 1.1 Why a first-principles frame is worth the effort

The case for first-principles models is not aesthetic. It is that they generalize out of distribution. A thermodynamic occupancy model parameterized on one promoter library predicts new promoters because the underlying physics is the same [12]; a deep learning model trained on the same library predicts new promoters only insofar as they resemble the training set. The contrast is not absolute: models trained on large libraries of random sequence do extrapolate to native promoters [70], and occupancy models fail out of distribution when their parameters depend on context. For the non-coding genome, where the relevant variation is by definition rare and the regulatory grammars are species-and tissue-specific [2], this distinction matters. The ground-up frame is the one that admits perturbative, mechanistic, and synthetic interrogation. We do not claim it is sufficient, Section 10 makes clear that it is not, but we claim it is necessary, and that the layered decomposition makes its sufficiency conditions explicit.

That argument is not weakened by the arrival of genome-scale prediction resources, but what it costs to check it has changed. Since September 2026 a precomputed catalogue of predicted molecular consequences for every possible single-nucleotide substitution in the human genome has been publicly available [55], which means the perturbation surface over which a layered decomposition makes its claims is, for the first time, a shared object rather than a per-laboratory expense. We treat this addition in Section 8, keeping the first-principles rationale above intact and asking a narrower question of the new technology: which of this paper’s predictions does it render resolvable, and which does it leave exactly where they were.

## 2. Layer 1: Sequence as Formal Language

The first layer asks: what kind of grammar does DNA require? Formal language theory provides a hierarchy of grammars (regular, context-free, context-sensitive, recursively enumerable) distinguished by the memory and structure their parsers need [7]. Mapping genomic features onto this hierarchy clarifies which biological objects can be recognized by simple machinery and which require more. The exercise is most useful when it predicts where finite-state methods will succeed and where they will fail.

### 2.1 The Chomsky hierarchy as a working hypothesis

Regular grammars suffice for memoryless local patterns: TATA boxes, polyadenylation signals, simple tandem repeats, and single transcription-factor binding motifs. These can be recognized by a finite-state automaton scanning a position-weight matrix, and most current motif-discovery tools are exactly that, sometimes with extensions. Much single-site transcription-factor recognition is well described at this level by position weight matrices [71], although motif spacing and orientation effects (“soft syntax”) go beyond it [72]. This is fortunate, because finite-state methods are computationally cheap and the corresponding biology (a protein domain reading a few base pairs) is physically minimal.

Context-free grammars (CFGs) become necessary when the sequence encodes nested dependencies. Secondary structure in RNA is the canonical example: stems require base i to pair with base j, and pseudoknot-free folds correspond exactly to derivations in a CFG. Stochastic CFGs (SCFGs), which assign probabilities to production rules, have been used productively for ncRNA structure prediction [13], tRNA modeling, and ribozyme classification [14]. The Rfam 15 database (2025) continues to use Infernal SCFG covariance models as its core annotation engine, with Rfamseq expanded by 76% to 26,106 genomes and 65 families enhanced with three-dimensional structural data [15]. SCFGs remain indispensable where sequence conservation is weak but structural conservation is strong, because covariation across paired positions can be inferred from the grammar.

However, the hegemony of hand-designed SCFGs is now challenged on two fronts. First, Onokpasa, Wild, and Wong (2024) demonstrated that automatically discovered grammars outperform human-expert SCFGs for joint sequence-structure compression of RNA families [16]. Second, deep-learning alternatives such as RNAformer outperform earlier single-sequence methods (F1 0.743 versus 0.714 for SPOT-RNA on the TS1 benchmark) and perform on par with the alignment-based SPOT-RNA2, without requiring multiple sequence alignments [17]. These results suggest that the formal-language frame retains value for interpretability and auditable reasoning but is no longer the performance frontier for pure prediction.

Context-sensitive grammars (CSGs) are invoked for chromatin-dependent regulation, enhancer–promoter looping, and splicing decisions whose outcome depends on flanking elements [8]. Here the formal claim should be made carefully. CSGs are appealed to because the relevant biology is genuinely context-dependent, but few published models are context-sensitive in the formal sense; most are CFGs with auxiliary feature inputs, or probabilistic graphical models that happen to capture the same dependencies more tractably. In practice, mammalian enhancer logic appears to live above the context-free level, but our parsers for it are not yet on the formal hierarchy.

### 2.2 Where the linguistic analogy is strongest, and where it leaks

Two empirical observations support the linguistic frame. First, non-coding sequence shows Zipf-like rank-frequency distributions of k-mers more clearly than coding sequence [18], consistent with a layered vocabulary of motifs of varying frequency, although Zipf-like distributions also arise in sequences with no linguistic structure, so this observation is weak evidence on its own [73,74]. Second, redundancy, measured as block entropy below the i.i.d. baseline, is higher in non-coding regions, which is what one would expect if regulatory signals are encoded with error tolerance against thermal noise and mutation [18]. These are not proofs that DNA is a language; they are observations that the statistical signature of a structured code is present.

The analogy leaks at three points. RNA tertiary structure (pseudoknots, kissing loops) cannot be expressed by a CFG and requires either crossing dependencies or auxiliary tape, putting it formally beyond context-free. Chromatin-mediated regulation operates on a three-dimensional substrate where adjacency is not linear, so the alphabet on which the “grammar” acts is not the genome read 5′→3′ but the contact graph. And combinatorial transcription factor logic at mammalian enhancers often depends on the precise stoichiometry of bound proteins rather than the sequence per se, which moves the relevant computation off the DNA tape and into the protein layer.

### 2.3 Stochastic CFGs as a concrete deliverable

Of the formal-language tools, SCFGs remain the one with the cleanest computable path from sequence to function. The pipeline, sequence → SCFG parser → maximum-likelihood structure → predicted function, is fully algorithmic and does not require labeled training data [13]. SCFGs handle gaps and unknown nucleotides probabilistically, which makes them robust for the deep alignments where conservation is loose. They are the layer-1 method most likely to give a domain expert something actionable: a sequence annotated with a structural prior whose grammar is auditable, not a black-box embedding. The practical recommendation is that SCFGs should be used for interpretable structural annotation, while deep models such as RNAformer should be used when raw predictive accuracy is the goal.

## 3. Layer 2: Information-Theoretic Structure of the Sequence

The second layer asks: independent of any specific grammar, what does the sequence look like as an information-bearing string? Three frameworks give partially overlapping answers, algorithmic information theory through compression, digital signal processing through spectra, and fractal geometry through DNA walks. Each picks up a different correlate of function, and each has known approximation pathologies that the literature too often glosses.

### 3.1 Compressibility as a proxy for algorithmic structure

Kolmogorov complexity K(x), the length of the shortest program that outputs x, is the canonical measure of algorithmic information [9]. It is also formally uncomputable, which means every empirical use is an approximation. In practice this means Lempel–Ziv complexity [20] or block decomposition methods on top of small reference universal Turing machines [21]. These are upper bounds on K, tight in some sequence regimes and loose in others.

A critical development in this area is the demonstration by Abrahão, Zenil, and colleagues (2024–2025) that the assembly index used in assembly theory is formally equivalent to a context-free-grammar Lempel–Ziv compression scheme [22,23]. This result has direct bearing on any claim that algorithmic complexity differentiates living from non-living molecular structure: the assembly index cannot perform better than Shannon entropy in stochastic scenarios and does not, by itself, detect the action of selection. The implication for our framework is that compressibility-based descriptors, while empirically useful, should be interpreted as measuring statistical regularity rather than genuine algorithmic structure.

With those caveats, complexity indices computed in this way are informative about nucleosome occupancy and can complement composition-based occupancy models [21]. The strongest defensible claim is that compressibility could provide a training-free, label-free annotation track; whether it captures algorithmic structure or acts as a proxy for composition and biophysical features remains open, and Section 9 tests this directly.

### 3.2 Spectral analysis: real signals and false positives

Mapping nucleotides to numerical values (Voss indicators, electron-ion potentials, or optimized spectral envelopes) and applying the discrete Fourier transform exposes periodicities [24]. Two are robust and biologically interpretable: the 3-bp periodicity from codon bias, which reliably marks coding regions independent of any database [25], and a ∼10.5-bp periodicity in some nucleosome-positioning contexts, which reflects the helical pitch of B-form DNA. Beyond these, spectral peaks at biologically named frequencies require care: short windows produce broad spectral leakage, and the choice of nucleotide-to-number mapping can manufacture spurious harmonics.

### 3.3 DNA walks and fractal dimension

DNA walks convert a sequence into a path in a metric space, typically by mapping purines and pyrimidines to step directions, and characterize the path geometrically [26]. The Hurst exponent H of such a walk distinguishes uncorrelated sequence (H ≈ 0.5, Brownian) from sequences with long-range correlations (H > 0.5, persistent). Intron-containing genes and many non-coding regions show H > 0.5; pure coding sequence is closer to H ≈ 0.5 [19]. The literature linking walk fractal dimension and lacunarity to disease-associated sequence alterations [27] should be read as suggestive rather than mechanistic.

### 3.4 The information-theoretic layer as an annotation stack

Taken together, the three frameworks of this layer are best understood as complementary annotation channels rather than competing theories. Compressibility flags algorithmic regularity; spectra flag periodic structure; DNA walks flag long-range correlation. None individually predicts function; all three together provide a low-cost, label-free prior that the higher layers can condition on. The interface to layer 3 is exactly this: information-theoretic descriptors mark where on the genome the runtime, mobile elements and repeat arrays, has been most active.

## 4. Layer 3: Mobile Elements as the Genome’s Runtime

The third layer is where the operating-system metaphor for non-coding DNA earns or loses its keep. Transposable elements (TEs) and tandem-repeat arrays make up roughly half the human genome by mass [28] and are routinely described in computational language: LINE-1 as a “copy-paste” subroutine, Alu as a parasitic macro hijacking the L1 reverse transcriptase, satellite DNA as structural padding, TAD boundaries as memory partitions.

### 4.1 Retrotransposons as algorithmic operators

A full-length, active LINE-1 element encodes ORF1p (an RNA-binding chaperone) and ORF2p (an endonuclease and reverse transcriptase), assembles into a ribonucleoprotein particle with its own transcript, and inserts a copy at a new genomic site via target-primed reverse transcription [29]. The insertion is not random: ORF2p prefers the cleavage motif 5′-TTTT/AA-3′ and shows strong dependence on local DNA structure. From the layered-stack perspective, an active L1 implements a sequence-conditioned write to a new memory address, with the sequence of the host DNA selecting the address.

Solovyov and colleagues (2025) published a pan-cancer multi-omic model of LINE-1 activity revealing locus-level heterogeneity of retrotransposition efficiency [30]. This work demonstrates that L1 elements function not as uniform copy-paste operators but as locus-specific agents whose activity is modulated by local chromatin state, DNA methylation, and the host piRNA defense system. Recent studies document L1 as both an oncogenic driver and an immunogenic target in cancer [31,75], and reverse-transcriptase inhibitors have been reported to induce autophagy in an ORF1p-dependent manner [76]. Zueva (2025) reviews the “plan or pandemonium” paradox of retrotransposon reactivation in cancer [32].

### 4.2 What TEs do to the regulatory layer

Three mechanisms are well-attested. First, promoter and enhancer co-option: a TE insertion brings its own regulatory sequences into a new neighborhood, and selection occasionally retains the resulting expression change [33]. Many lineage-specific regulatory innovations in mammals trace to this mechanism. Second, chromatin architecture: certain TE families, notably SINEs, are enriched at TAD boundaries [77], and some TE-derived sequences act as insulators [34], suggesting that the spread of these elements has shaped the spatial organization of the genome. Third, exonization: an intronic TE can be spliced into a host transcript, contributing protein-coding or regulatory sequence depending on its content.

### 4.3 Satellite DNA and structural delimiters

Tandem-repeat arrays at centromeres and telomeres have algorithmic complexity close to the lower bound, they are nearly maximally compressible, and that low complexity is the functional signal. Pericentromeric heterochromatin, telomere maintenance, and chromosomal segregation depend on the highly repetitive character of these arrays rather than on any specific sequence. Tools such as TAREAN [35] and SatXplor [36] reconstruct consensus monomers from k-mer abundance distributions, an approach that works precisely because the underlying algorithmic structure is “repeat this short string.”

### 4.4 TE dynamics as a population-level automaton

The aggregate dynamics of a TE family in a host genome can be modeled as a competition between transposition rate, selection coefficient, and host repression. Population-genetic models of the first steps of invasion show that elements with regulated transposition can invade and then stabilize, whereas elements with a constant rate are usually lost to drift or sterilize their host [37]. Such models suggest that the “junk-DNA landscape” reflects a balance between transposition, regulation and selection rather than the residue of neutral drift alone.

## 5. Layer 4: Compilation from Sequence to Interaction Networks

The fourth layer is where the layered frame is at its strongest and most testable. Compilation here means converting static sequence into the dynamic interaction networks that determine cellular state. Two formalisms do most of the work: thermodynamic occupancy modeling, which converts promoter sequence into a binding-state probability distribution [10], and rule-based modeling, which converts protein-domain logic into a graph-rewriting simulation [38]. Together they take us from layer-3 outputs (which sequences exist where) to layer-5 inputs (what regulatory dynamics those sequences produce).

### 5.1 Thermodynamic occupancy as the prokaryotic gold standard

For prokaryotic and simple eukaryotic promoters, the thermodynamic occupancy model is the cleanest mechanistic theory we have. The promoter is treated as a statistical-mechanical ensemble of binding configurations: free DNA, RNA polymerase bound, individual transcription factors bound, cooperative complexes bound. Each configuration s has a free energy ΔG(s) computed from the sequence via energy matrices. The occupancy probability of a configuration is its Boltzmann weight divided by the partition function Z = Σ exp(−ΔG(s)/kBT) over all configurations, and gene expression is taken as proportional to the occupancy of transcriptionally productive states.

The model derives logic-gate behavior from physics rather than postulating it. Two cooperatively binding activators implement a soft AND gate; a repressor whose binding site overlaps the polymerase footprint implements a NOT gate; a competing-binding pair gives an XOR-like response when stoichiometries are tuned [39]. Sort-Seq and related massively parallel reporter assays, combined with Bayesian or MCMC inference, recover energy matrices accurate to roughly 1 kBT per base on well-behaved promoters [12]. At that resolution, the model predicts new promoter behavior quantitatively, not just qualitatively.

A caveat: this works cleanly for prokaryotic promoters and for simple, isolated eukaryotic enhancers in cell-free or reporter systems. Occupancy-style models have been extended to *Drosophila* enhancers [78] and to yeast promoter libraries [70], but for native mammalian enhancers, where the relevant logic involves chromatin state, cooperative protein assemblies of unknown stoichiometry, and looping to distal targets, thermodynamic occupancy models become considerably less predictive.

### 5.2 Rule-based modeling above the partition function

Once individual binding interactions are quantified, the next problem is combinatorial. A protein with five phosphorylation sites can occupy 32 internal states; a heterodimer of two such proteins has more than 1,000. Rule-based modeling languages, Kappa [38] and BioNetGen [40] are the two mature platforms, sidestep this by representing molecules as structured graph objects with named binding sites and modification states, and representing interactions as local graph-rewriting rules.

A significant 2024 advance is the release of MCell4 with BioNetGen integration via a new open-source library, libBNG, which provides the capability to run BioNetGen rule-based models in a spatially resolved Monte Carlo simulation environment through a Python API [41]. This integration bridges the gap between well-mixed and spatially extended rule-based modeling, enabling the first production-ready spatially resolved stochastic simulations of rule-based signaling networks at the subcellular scale.

## 6. Layer 5: Execution as Cellular and Multicellular Dynamics

The fifth layer is where the compiled regulatory networks are actually run. Three formalisms cover most of what is currently tractable: membrane computing for compartmentalized intracellular dynamics, neural cellular automata for tissue-scale morphogenesis, and whole-cell models for the integrative limit.

### 6.1 Membrane computing and compartmental rules

P-systems [79] abstract a cell as a hierarchy of nested membranes, each containing a multiset of objects (molecules) and a local rule set (reactions). A widely cited result is that P-systems are Turing-complete. This is sometimes invoked as evidence that cells “are universal computers.” It is worth being careful here: Turing-completeness is a low bar, achieved by many trivial systems, and what matters for biology is not whether a P-system can in principle compute any function but whether the specific rule sets that correspond to actual cellular biochemistry produce the observed behavior.

### 6.2 Neural cellular automata and morphogenesis

Neural cellular automata (NCAs) [80] model a tissue as a grid of cells running an identical update rule that depends on local neighborhood state. The shared rule is the analogue of the shared genome; the global pattern that emerges is the phenotype. A comprehensive synthesis by Hartl, Levin, and Pio-Lopez (2025) surveys NCA applications across molecular, tissue, and organismal scales, including evolvable update rules and their relationship to biological development [42]. Recent results include Béna and colleagues’ work on universal neural cellular automata at GECCO 2025 [43], demonstrating that a single NCA architecture can learn multiple developmental programs; Kvalsund and colleagues’ active-sensing NCAs [81]; Chow and Bentley’s finding that an NCA model of morphogenesis develops early, generalized factors analogous to evolutionarily conserved transcription factors and conserves them when learning new body forms [82]; and Guichard and colleagues’ ARC-NCA applying NCAs to abstraction-and-reasoning tasks at ALIFE 2025 [83].

The ENIGMA family of models structures the update rule as an explicit gene regulatory network, with simulated genes regulating each other and producing signaling molecules whose gradients drive cell behavior [44]. These models reproduce homeotic transformations, pattern maintenance across tissue sizes and simple regulatory motifs seen in development [44]. If compact gene regulatory networks suffice to specify such multicellular geometries, the algorithmic information required to specify a developmental program is smaller than naive counting of cell types and positions would suggest.

### 6.3 Whole-cell models and the integrative limit

Whole-cell models attempt to simulate every molecular species and reaction in a cell over a complete cell cycle. The pioneering Karr et al. (2012) model of *Mycoplasma genitalium* [45] integrated 28 sub-models covering 525 genes. Subsequent efforts have extended the approach to *Escherichia coli*, where the Covert Lab’s whole-cell model accounts for the functions of 43% of characterized genes [46,84]. Sun, DeFelice, Gillies, and colleagues (2024) used the *E. coli* whole-cell model to cross-evaluate operon structures genome-wide, demonstrating that the model can predict the phenotypic consequences of gene rearrangements that have not been experimentally tested [47]. Karr and colleagues have proposed a human whole-cell modeling effort that uses H1 human embryonic stem cells as its initial testbed [85], a first step toward mammalian whole-cell simulation.

## 7. Synthetic Validation: Does the Stack Compose?

### 7.1 Cello and CELLM: thermodynamics-aware compilation to DNA

Cello is a CAD tool that takes a Verilog specification of a logic circuit and produces a DNA sequence implementing it [48]. The pipeline maps each Verilog gate to a characterized DNA “gate”, a promoter– repressor pair, and uses thermodynamic models to ensure that the output flux of one gate is compatible with the input threshold of the next. Cello-designed circuits routinely operate as specified in *E. coli* with high success rates [49]. As a test of the layered frame, Cello succeeds because it stays within the regimes where the underlying physics is well-characterized.

A notable 2025 development is CELLM, developed by Abello Castillo and Gutiérrez Pescarmona, which integrates Cello 2.1 with large language models (DeepSeek-R1, Phi-4) through LangChain to translate natural-language circuit descriptions into DNA designs [50]. CELLM represents the convergence of two previously separate paradigms, formal-language-based genetic circuit design and neural-network-based text understanding, and shows that circuit compilation can be driven from natural-language specifications. Machine-guided design of cell-type-specific enhancers [86,87] extends compilation to eukaryotic regulatory sequence, but with data-driven rather than thermodynamic models.

### 7.2 Gro: programming multicellularity

Gro is a programming language and simulator for multicellular behavior, allowing a researcher to specify colony growth, cell–cell signaling, and division dynamics in high-level terms and observe the resulting spatial patterns [51]. Its predictions match experimental colony patterns reasonably well in well-studied systems. As with Cello, the success rate degrades as the target organism’s biology departs from the parameters that the simulator assumes.

### 7.3 Foundation models as oracles for layer 1

Genomic foundation models such as Evo 2 are large autoregressive language models trained on the full sequence diversity of life [5]. They are not ground-up models in the sense of this paper, they are data-driven, but they are informative for the layered frame in two ways. First, they provide a benchmark: any first-principles theory of regulatory grammar should at minimum reproduce what a foundation model recovers from raw sequence, and where it does not, that is a place where the first-principles theory is incomplete. Second, the internal representations of these models, when probed via sparse autoencoders [6], recover known biological grammars without explicit annotation, which is consistent with the linguistic frame of layer 1. Evo 2’s 40-billion-parameter StripedHyena 2 architecture with 1-megabase context represents a qualitative step beyond the earlier Evo 1 model [52], and its ability to generate synthetic eukaryotic genome sequences provides a new validation paradigm for the layered stack. The release of a precomputed, genome-wide variant-effect catalogue changes what an oracle of this kind can be used for, and we treat that development separately in Section 8.

## 8. AlphaGenome Atlas: A Precomputed Perturbation Surface for the Layered Stack

Section 7.3 treated genomic foundation models as oracles for layer 1. On 8 September 2026 that relationship changed in kind rather than in degree, with the release of AlphaGenome Atlas by Google DeepMind [55]. The rationale set out in Sections 1 and 1.1 is unaltered: the case for first-principles models still rests on out-of-distribution generalization, and a data-driven oracle cannot supply that, however large it becomes. What the Atlas changes is the cost and the coverage of testing the layered decomposition. The perturbation surface over which every prediction of this paper is stated, what happens to molecular phenotype when a given base changes, has become a public, precomputed object rather than a per-laboratory expense. This section sets out what that makes newly resolvable at each layer, and what it structurally cannot supply.

### 8.1 What the resource contains

AlphaGenome Atlas is a one-petabyte database of precomputed AlphaGenome predictions for every possible single-nucleotide variant in the human reference genome, approximately nine billion substitutions, together with predictions for many observed insertions and deletions [55,59]. For calibration, this is more than thirty times the size of the AlphaFold Database [59]. The underlying model takes one megabase of sequence as input and predicts thousands of genome tracks across eleven output types, RNA-seq expression, CAGE and PRO-cap transcription initiation, splice sites, splice-site usage, splice junctions, DNase and ATAC accessibility, histone modifications, transcription-factor binding, and chromatin contact maps, over hundreds of human and mouse cell types and tissues [56].

Three derived layers matter for the argument of this paper. First, the AlphaGenome Variant Impact (AVI) score, a single scalar per variant that combines AlphaGenome’s regulatory predictions with AlphaMissense’s protein-level predictions [57] and with conservation and variant-consequence features, so that coding and non-coding variants are ranked on one scale; the authors report best-in-class performance across many variant-pathogenicity and rare-disease benchmarks [55]. Second, AVI feature attributions, which decompose each score additively into interpretable categories such as chromatin accessibility, splicing, expression, conservation, and protein impact. Third, a genome-wide compendium of more than 2,500 recurrent cis-regulatory motifs together with their locations. The resource is free for non-commercial use through a web portal that requires no coding, through the AlphaGenome API, and as a skill within Google’s Antigravity agentic development platform [55,59].

### 8.2 Why this extends the rationale rather than replacing it

The Atlas is a data-driven artifact and its contents are predictions, not measurements. Both DeepMind’s own framing and independent coverage of the release are explicit on this point: the resource is a research aid that has not been validated for clinical use, and its predictions are forecasts of what could happen if a letter changed rather than observations of what did [55,58,59]. The first-principles case stands where it stood, and Section 7.3’s characterization of foundation models as oracles rather than theories stands with it.

The addition is methodological, and it is substantial. A layered decomposition makes a claim about propagation: a base substitution alters a string (layer 1), alters its information-theoretic descriptors (layer 2) within a context set by mobile-element history (layer 3), shifts an occupancy or rule-firing distribution (layer 4), and produces a change in cellular dynamics (layer 5). Testing such a claim requires a dense perturbation surface, ideally the complete counterfactual: for every base, what happens if it changes. Until now that object did not exist. In-silico saturation mutagenesis was available locus by locus at the cost of a model run per variant, which meant every test of the framework began with a discretionary choice of loci and inherited whatever bias that choice carried. The Atlas removes the choice by exhausting the single-substitution space. For the first time the layered frame can be evaluated with the genome as the denominator.

The second gain is that the Atlas is modality-resolved. A single scalar impact score would be of limited use here, because the layered claim is not that a variant matters but that it matters **through a particular layer**. The AVI feature attributions and the underlying per-track predictions expose exactly that: a variant can be labeled as acting predominantly through splicing, accessibility, expression, or protein sequence. This is a machine-readable form of the layer assignment that Sections 2 through 6 assert on theoretical grounds, which means the assignment can now be audited at scale rather than argued case by case.

### 8.3 What becomes newly testable, layer by layer

#### Layer 1 (sequence as formal language)

The motif compendium is an empirically derived, positioned lexicon of the “words” that Section 2 posits, and it converts Table 1 from a taxonomy into a measurable partition. For any variant with a substantial predicted effect one can ask what the minimal grammar class needed to account for that effect is: disruption of a single local motif is a regular-language event recognizable by a finite-state scan; an effect that requires a paired dependency, donor with acceptor, or a base pair in a folded RNA, is context-free; an effect that appears only when a distal partner and a chromatin state are jointly specified sits above the context-free level, and the model’s contact-map channel makes the distal partner explicit. The fraction of consequential variants falling in each class becomes a computable quantity rather than an assertion, and the claim in Section 2.1 that a large fraction of cis-regulatory recognition is well captured at the finite-state level acquires a denominator. An early use of the resource by Zeitlinger and Weilert, separating transcription factors that only alter accessibility from those that also alter expression [55], is precisely this kind of operational layer assignment.

**Table 1.**
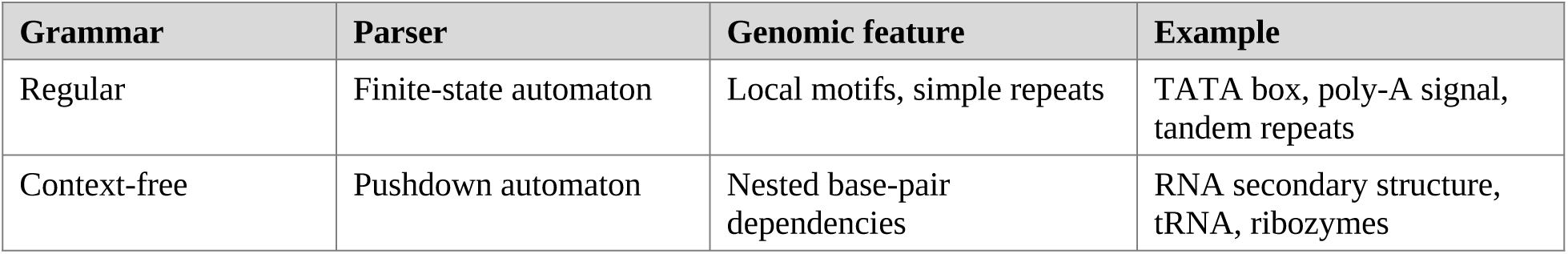
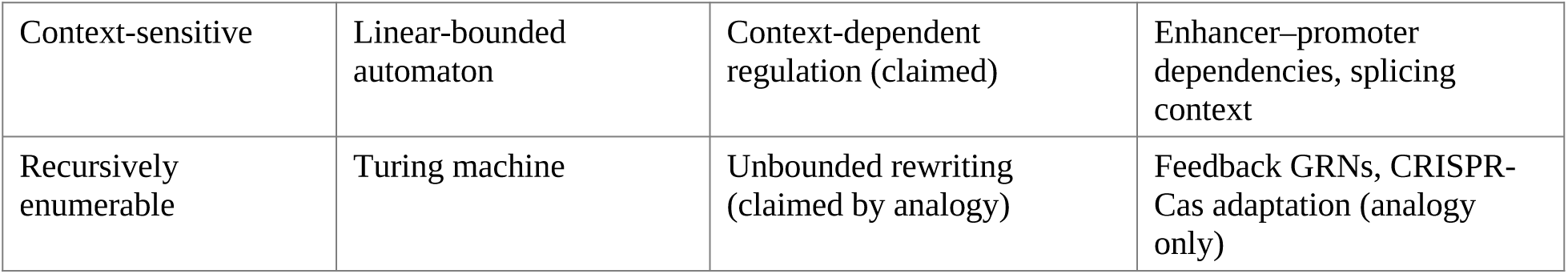
Chomsky hierarchy applied to genomic elements. Each grammar class corresponds to a parser of distinct computational power; the genomic features at each level reflect this distinction. The last row is an analogy: feedback networks are dynamical systems rather than string recognizers.

| **Grammar** | **Parser** | **Genomic feature** | **Example** |
|---|---|---|---|
| Regular | Finite-state automaton | Local motifs, simple repeats | TATA box, poly-A signal, tandem repeats |
| Context-free | Pushdown automaton | Nested base-pair dependencies | RNA secondary structure, tRNA, ribozymes |
| Context-sensitive | Linear-bounded automaton | Context-dependent regulation (claimed) | Enhancer–promoter dependencies, splicing context |
| Recursively enumerable | Turing machine | Unbounded rewriting (claimed by analogy) | Feedback GRNs, CRISPR-Cas adaptation (analogy only) |

#### Layer 2 (information-theoretic structure)

Prediction (1) of Section 11 has been difficult to test because it requires a dense regulatory-potential target that is not itself derived from k-mer composition, and a conservation covariate measured on the same footing. The Atlas supplies a continuous target at every base, and conservation appears as one of the AVI attribution categories, so the covariate the prediction most needs to be partialed out arrives in the same table as the response. The regression, algorithmic-complexity tracks as predictors, k-mer composition, GC content and conservation as covariates, can now be run genome-wide on public data, without new experiments and without a curated locus set. A first test of the prediction that does not depend on the Atlas, using measured rather than predicted effects, is reported in Section 9.

#### Layer 3 (mobile elements as runtime)

Because coverage is genome-wide, the Atlas necessarily covers the roughly half of the genome that is transposable-element derived. Per-family and per-age distributions of predicted impact give a quantitative measure of how much regulatory load each mobile-element family carries, and the strong form of the co-option claim in Section 4.2 becomes checkable: co-opted copies should show impact distributions shifted relative to matched non-co-opted copies of the same family at similar sequence divergence. One caveat is specific to this layer. The training data are mapping-biased in repetitive sequence, so model behavior inside young, high-identity elements is the least reliable part of the resource and should be treated as such rather than as a null result.

#### Layer 4 (compilation)

This is where the Atlas bears most directly on the framework, because a thermodynamic occupancy model of a regulatory element implies a specific structure for that element’s complete single-base mutational surface: additive contributions from independent sites, plus deviations wherever binding is cooperative. The Atlas provides that surface for every promoter and every candidate cis-regulatory element in the genome. An energy-matrix model can be fitted on a subset of bases and tested for extrapolation to held-out bases and held-out loci, and, more informatively, its residuals can be examined for where the physics departs from the prediction. The expected pattern, promoters well predicted and distal enhancers poorly, coincides with the Atlas’s own weakest domain, which is itself the useful observation: a first-principles occupancy model and a trained sequence model degrade in the same place, which points at a missing ingredient, chromatin state and three-dimensional context, rather than at insufficient parameters or insufficient training data.

#### Layer 5 (execution)

The Atlas stops at molecular phenotype. It predicts tracks, not cell-state trajectories, morphogenesis, or organismal traits. The population-genetic application reported at release is instructive on exactly this boundary: grouping rare non-coding variants by predicted molecular effect across whole-genome data from more than 54,000 UK Biobank participants recovered 22% more independent non-coding associations than the same analysis without Atlas filtering, and restricting attention to the 1% of variants predicted most impactful identified nineteen genomic regions associated with body mass index [55,59]. That is a molecular-to-organismal relay accomplished statistically. The layered claim is that the same relay can be made mechanistically, through layers 4 and 5, and the composability problem of Section 10.6 now has a well-posed form: given a molecular effect that the Atlas predicts and orthogonal data confirm, do rule-based and gene-regulated cellular-automaton models of the relevant network reproduce the cellular consequence?

### 8.4 What the Atlas concedes to the layered hypothesis

The limits of the resource are as informative as its coverage, because several of them are structural rather than provisional, and they fall where the layered frame claims to be necessary.

#### Combinatorics

Nine billion is exhaustive for single substitutions and nil for combinations. The number of variant pairs within a single 500-bp regulatory element already exceeds a million, and the genome-wide pair space is of order 10¹⁹, permanently beyond precomputation, at any plausible compute budget. Epistasis between sites in the same element is exactly what a partition function over binding configurations represents natively (Section 5.1) and what an additive attribution scheme cannot represent at all. The Atlas’s own interpretability layer is additive by construction, so cooperativity is the one thing its decomposition is structurally unable to express. This is the clearest concession the new technology makes to the argument of this paper: the interaction surface cannot be precomputed. It has to be computed on demand, either by running a sequence model such as AlphaGenome on each combination or by a mechanistic model, and the layered claim is that the mechanistic model will do better where binding is cooperative.

#### Locality

Distal regulatory effects and long-range interactions remain the model’s weakest domain, as its authors state [56]. Section 10.5 makes the parallel observation about energy models. The coincidence is the substantive point, not an embarrassment for either method.

#### Modality ceiling

The output space is the eleven track types the model was trained to predict. Anything not represented there, post-translational modification, protein stoichiometry, metabolic state, mechanical context, is invisible by construction, and Section 2.2 already identified stoichiometry-dependent enhancer logic as a place where the relevant computation leaves the DNA tape. A variant predicted null is null only with respect to the modalities modeled.

#### Reference frame

Variants are scored one at a time against the reference genome. Real genomes carry them in haplotype context, alongside structural variation, somatic mosaicism, and an epigenetic state with its own history, the same set of factors that Section 10.6 identifies as the reason composition across layers is only partial.

#### Circularity

The most important methodological constraint is that a layered model validated against Atlas predictions has been tested for agreement with another model, not with nature. Used well, the resource is a screen and a matched-control generator: it makes it cheap to assemble large, well-matched sets of predicted-high-impact and predicted-null variants, which is precisely what the falsification conditions of Section 11 require. Adjudication stays with experiment and orthogonal measurement, massively parallel reporter assays, base-editing screens, QTL data, and case resolution of the kind reported for a DNM1 variant at release, where the predicted mechanism, a created splice site producing an abnormally extended protein, was confirmed by experimental screens [55].

### 8.5 How the hypothesis becomes resolvable

Table 3 restates each of the five predictions of Section 11 in terms of what the Atlas now makes feasible. The pattern is consistent: the predictions that were bottlenecked on the absence of a dense, unbiased, modality-resolved perturbation surface are now executable on public data, while those that depend on interaction structure or on execution-layer dynamics are not, and remain the province of the ground-up formalisms. We have not run the genome-wide analyses here; Section 9 reports the one test we did run, on measured rather than predicted effects.

**Table 2.** Mobile and repetitive elements as computational primitives.

| **Element** | **Computational role** | **Mechanism** | **Functional consequence** |
|---|---|---|---|
| LINE-1 | Subroutine with state | Target-primed reverse transcription | Insertional mutagenesis; promoter creation; copy-paste write |
| Alu / SINE | Macro / mobile module | Trans-complementation by L1 machinery | Splice-site donation; mRNA stability change; chromatin boundary |
| Satellite DNA | Structural delimiter | Tandem repetition (low K complexity) | Centromere integrity; heterochromatin compaction |
| TAD boundary | Memory partition | CTCF / cohesin loops | Insulates regulatory domains; prevents enhancer crosstalk |

**Table 3.** The five predictions of Section 11 restated against what AlphaGenome Atlas makes feasible. The resource resolves the sampling and coverage bottleneck; it does not resolve the interaction or execution bottlenecks.

| **Prediction (Section 11)** | **What was missing** | **Atlas-enabled test** | **Status** |
|---|---|---|---|
| (1) Compressibility predicts regulatory activity independently of sequence content | A dense regulatory-potential target not derived from k-mer composition, plus a matched conservation covariate | Genome-wide regression of LZ/BDM tracks on AVI and per-modality effect magnitude, controlling for k-mer composition, GC and the conservation attribution | Executable now on public data; first test on measured MPRA data (Section 9) does not support it at the variant level |
| (2) SCFG structural priors improve non-coding RNA variant classification | Enough labeled ncRNA variants at sufficient density to power a matched comparison | Saturation surfaces at ncRNA loci give a dense comparison baseline; SCFG-derived priors tested for added value over AVI alone | Executable now, with experimental labels still required for adjudication |
| (3) Thermodynamic occupancy extends predictively given measured parameters | Complete single-base mutational surfaces for many elements at once | Fit energy matrices on subsets of the Atlas surface; test extrapolation to held-out bases and loci; inspect residuals at promoters versus distal elements | Executable now; the informative result is where both methods fail together |
| (4) Rule-based reconstruction identifies the causal chain from variant to phenotype | A principled way to select which variants are worth reconstructing | AVI feature attributions nominate the entry layer (splicing, accessibility, expression) for each candidate variant, defining the input to the rule-based model | Partially enabled; layer-5 validation still required |
| (5) Small gene-regulated NCAs reproduce developmental robustness | A link from sequence perturbation to developmental dynamics | None directly; the Atlas terminates at molecular phenotype | Not addressed by the Atlas |

Two further predictions arise specifically from the new resource, and are stated here rather than in Section 11 because both are about the boundary between precomputation and mechanism.

(6) Grammar-class stratification. The minimal grammar class required to explain a variant’s predicted effect should track the modality of that effect. Splicing effects and promoter-proximal expression effects should be largely accounted for by local motif disruption, regular, with context-free extensions where paired splice signals are involved, whereas distal expression effects should require a jointly specified partner locus and chromatin context. Falsification: after matching on predicted effect size, local motif disruption accounts for distal cis-regulatory variant effects as well as it accounts for promoter-proximal ones, in which case the Chomsky-hierarchy stratification of Table 1 carries no empirical content.

(7) The epistasis gap. For pairs of variants within the same regulatory element, additive combination of single-variant Atlas scores, and AlphaGenome run directly on the double mutant, should systematically underperform a fitted thermodynamic-occupancy or rule-based model in predicting measured double-mutant effects, and the shortfall should grow with the cooperativity the occupancy model infers. Falsification: additive combination or direct model prediction matches measured double-mutant effects within experimental error across cooperativity regimes, in which case the layer-4 formalism adds nothing that data-driven prediction cannot supply, and the strongest remaining argument for the layered frame is lost.

The Atlas does not settle whether the genome is usefully treated as the input to a computation. It does something narrower and more useful: it makes the question expensive to dodge. Three of the five standing predictions can now be tested on public data at genome scale, and the two above address the region where precomputation cannot follow. If the layered decomposition is right, that region is exactly where its value should be visible.

## 9. A First Test of Prediction 1 on Measured Mutational Effects

Prediction (1) of Section 11 states that algorithmic-complexity descriptors should carry information about regulatory function beyond k-mer composition, GC content and conservation. Section 8.3 describes how the Atlas makes that regression executable genome-wide on predicted effects. The claim can, however, be tested first against measurements, which avoids the circularity set out in Section 8.4. Saturation-mutagenesis massively parallel reporter assays (MPRAs) supply exactly the perturbation surface the prediction needs: every possible substitution at every base of a regulatory element, each with a measured effect on reporter expression. We therefore tested a variant-level form of Prediction (1) on the disease-associated regulatory elements assayed by Kircher and colleagues [60]. The operational question is whether the algorithmic complexity of the sequence surrounding a base predicts how strongly mutating that base changes regulatory output.

### 9.1 Design

After removing duplicate genome-build entries, restricting to single-nucleotide substitutions and requiring at least ten barcodes per variant, the dataset comprised 38,556 variants in 29 element-by-condition datasets drawn from 21 regulatory elements at 20 distinct loci (Methods). The response was the absolute measured log₂ effect of each variant. Four descriptors of the hg38 reference sequence in a window centered on the mutated base were compared: Lempel–Ziv complexity (LZ76) [61], the Block Decomposition Method (BDM) estimate of algorithmic complexity [21,62], Shannon entropy and the zlib compression ratio. A 51 bp window was designated as primary, with 21 and 101 bp reported as sensitivity analyses. Every model included element fixed effects, so no estimate derives from differences in mean activity between elements, together with GC content, the first ten principal components of the 64 trinucleotide frequencies, and per-base phyloP100way conservation [63]. Standard errors were clustered by element.

### 9.2 In a pooled model, complexity is associated with effect size, and conservation does not absorb it

Successive terms raised the variance explained from 0.1441 (element alone) to 0.1451 (+GC), 0.1600 (+trinucleotide composition), 0.1689 (+conservation) and 0.1743 (+BDM) (Figure 2a). In the full model the BDM coefficient was −0.0303 per standard deviation (p = 0.0014): more compressible local sequence is associated with larger measured variant effects. Prediction (1) did not fix a sign, and this direction runs against the error-tolerance reading of Section 2.2, under which redundant sequence should buffer substitutions. The covariate most likely to explain this away did not. Without the conservation term the BDM coefficient was −0.0284 (p = 0.0022); with it, −0.0303, and the same held for every descriptor (Figure 2b). Complexity and conservation therefore carry separable information about these elements. A block-permutation null that circularly shifts BDM within each element, preserving its autocorrelation, gave a null coefficient of −0.00032 ± 0.00685 (s.d.); the observed value lay outside all 1,000 permutations.

**Figure 2.**
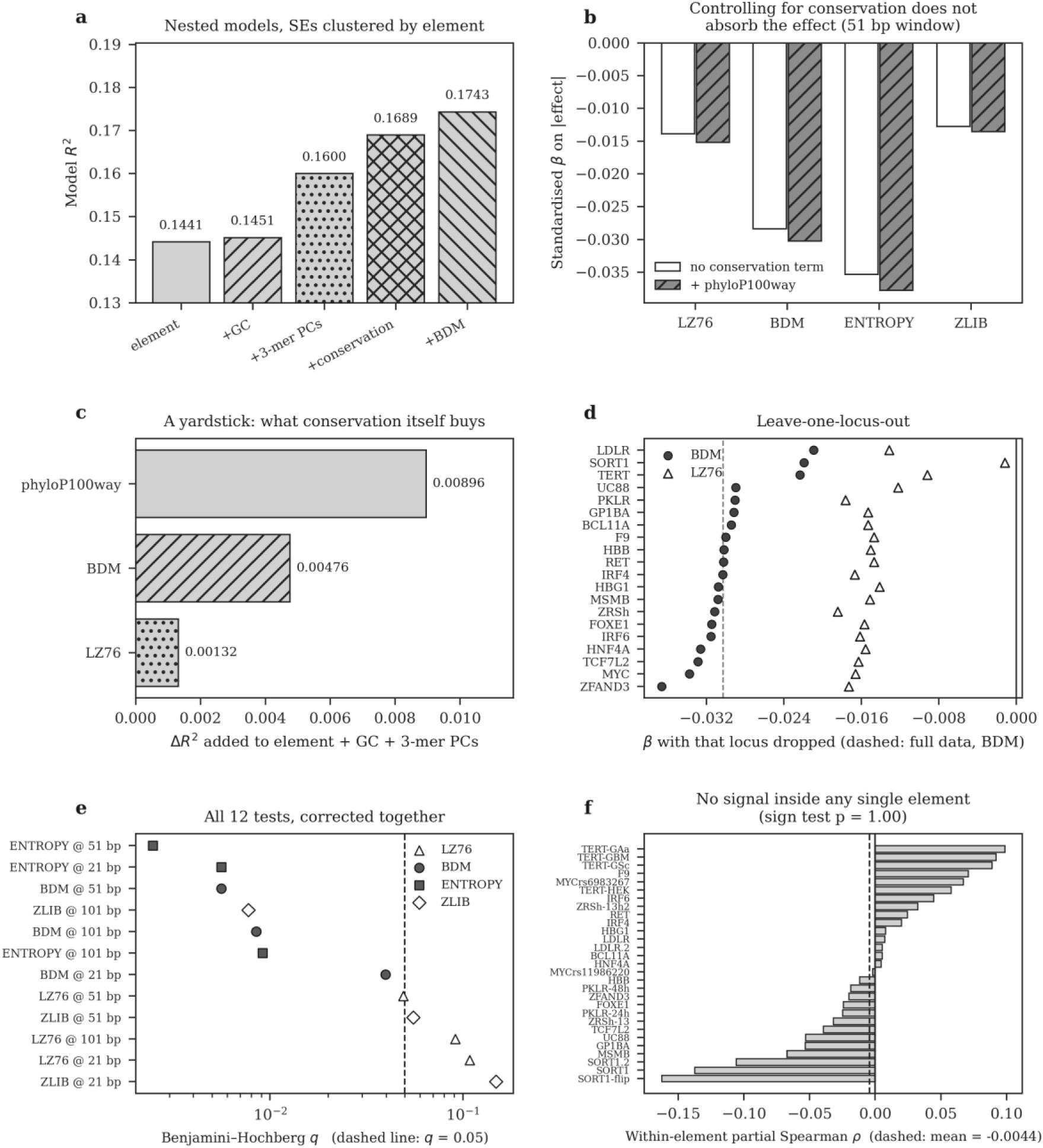
Algorithmic complexity of local sequence and regulatory variant effect size in saturation-mutagenesis MPRA (38,556 variants, 29 element datasets, 20 loci). (a) Variance explained by nested models of absolute variant effect; standard errors clustered by element. (b) Standardized coefficients for four complexity descriptors in the primary 51 bp window, without (open) and with (hatched) a per-base phyloP100way conservation term. (c) Increase in RZ over the composition-controlled model from conservation, BDM and LZ76, on a common scale. (d) Coefficients for BDM (filled circles) and LZ76 (open triangles) with each of the 20 loci removed in turn; dashed line, BDM estimate on all data. (e) Benjamini–Hochberg q values for all twelve descriptor-by-window tests; dashed line, q = 0.05. (f) Within-element partial Spearman correlations between BDM and absolute effect, controlling for GC, trinucleotide composition and conservation; dashed line, mean.

The effect is small, and the useful comparison is with conservation itself. Added to the composition-controlled model, phyloP100way raised RZ by 0.0090, BDM by 0.0048 and LZ76 by 0.0013 (Figure 2c). BDM complexity is thus a weak predictor of regulatory effect size, of roughly half the magnitude of per-base conservation, rather than a strong one.

### 9.3 Multiplicity and dependence on single loci

Four descriptors at three window sizes give twelve tests, which were corrected together by the Benjamini– Hochberg procedure [64]. All 12 coefficients were negative and 8 survived at q < 0.05 (Figure 2e). The primary test gave q = 0.0056; Shannon entropy was the strongest single descriptor (β = −0.0378, q = 0.0025), while LZ76 and zlib at 51 bp were borderline (q = 0.049 and 0.055). Because the SORT1 enhancer appears three times in the dataset (SORT1, SORT1-flip and SORT1.2), any property of that one locus enters the pooled estimate three times. Excluding it removed the LZ76 association entirely (β = −0.0012, p = 0.68) but not the BDM association (β = −0.0219, p = 0.0076). Dropping each of the 20 loci in turn, the BDM coefficient remained negative in every case, between −0.0366 and −0.0210 (Figure 2d). The LZ76 result should therefore not be counted as evidence for the prediction; the BDM and entropy results are tested further in Section 9.5.

### 9.4 The association is not present within individual elements

Residualizing both effect size and BDM on GC, trinucleotide composition and conservation separately inside each of the 29 element datasets, the partial Spearman correlation averaged −0.0044, was negative in only 14 of 29 elements (sign test p = 1.00; Wilcoxon signed-rank p = 0.90), and ranged from strongly negative at the SORT1 locus to positive at the TERT promoter (Figure 2f). The pooled estimate, which constrains covariate effects to be common across elements and weights large effects linearly, does not decompose into a rank association that holds inside each element. Complexity does not, on this evidence, identify which base within a given enhancer matters most, and a meta-analysis of per-locus slopes reaches the same conclusion (Section 9.5).

### 9.5 Robustness to composition, motifs, replication structure and measurement noise

We re-implemented the analysis from the public data and reproduced the reported estimates to within 0.0001 (BDM β = −0.0302 against −0.0303 reported, p = 0.0014; without SORT1, β = −0.0218 against −0.0219, p = 0.0077), then tested alternative explanations for it (Figure 3a; Methods). Unless stated otherwise, errors were clustered by dataset as in the primary model. Composition: replacing GC and ten trinucleotide components with mono-, di-and trinucleotide frequencies, including quadratic mononucleotide terms, roughly halved the coefficient to −0.016 (95% CI −0.025 to −0.007), and BDM measured relative to 50 dinucleotide-preserving shuffles of each window [88] gave −0.019 (95% CI −0.031 to −0.007). Shannon entropy, which depends on base composition alone, lost significance under full composition control (p = 0.11), and shuffle-adjusted LZ76 carried no signal (p = 0.60). Motifs: low-complexity windows overlapped more predicted transcription-factor sites [89], and motif-disruption covariates explained about five times more variance than BDM (ΔRZ = 0.026 versus 0.005); with them in the model the coefficient was −0.021 (95% CI −0.037 to −0.005).

**Figure 3.**
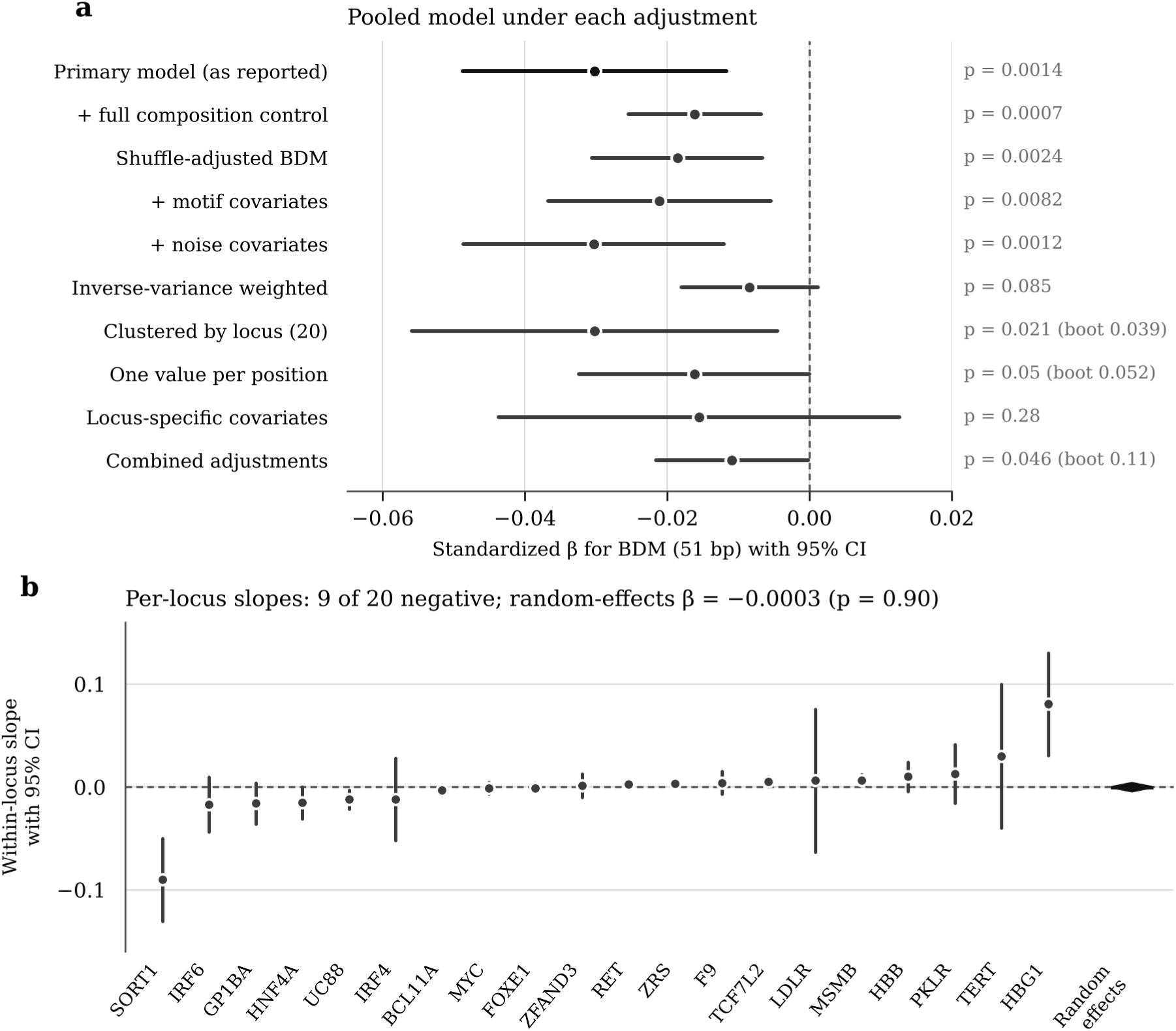
Robustness of the association between local compressibility (BDM, 51 bp window) and measured variant effect size. (a) Standardized coefficient with 95% confidence interval for the primary pooled model and for models that add full composition control (mono-, di-and trinucleotide frequencies), replace BDM by its excess over 50 dinucleotide-preserving shuffles, add motif-disruption covariates (JASPAR 2024 sites at p < 10⁻⁴) or measurement-noise covariates (log barcode count, standard error), weight by inverse variance, cluster errors by locus, collapse the substitutions and datasets at each base to one value per position, allow covariate effects to differ by locus, or combine shuffle-adjusted BDM, full composition control, and motif and noise covariates. Errors are clustered by dataset, as in the primary model, except in the last four rows, which cluster by locus. Bootstrap values (boot) are wild cluster bootstrap p values over 20 loci. (b) Within-locus slopes, with covariates fitted and BDM standardized separately for each locus, and their random-effects summary (diamond). Dashed lines, zero.

Replication structure: each window is shared by the three substitutions at a base, and six loci contribute several datasets. Clustering by locus rather than by dataset gave p = 0.021 (wild cluster bootstrap [90] p = 0.039), and collapsing to one value per position (9,834 positions), also clustered by locus, gave −0.016 (95% CI −0.032 to 0.000; bootstrap p = 0.052). Measurement noise: low-complexity windows had slightly fewer barcodes and noisier estimates (Spearman ρ between BDM and barcode count 0.10, and between BDM and standard error −0.13), and the absolute value of a noisy estimate is inflated by its noise. Adding barcode count and standard error as covariates left the estimate unchanged at −0.030 (95% CI −0.049 to −0.012), whereas inverse-variance weighting reduced it to −0.008 (95% CI −0.018 to 0.001); the association persisted for signal-to-noise (|z|, p = 0.006) and for significance calls (p = 0.011). Heterogeneity: allowing covariate effects to differ by locus, with errors clustered by locus, gave −0.016 (95% CI −0.044 to 0.013; p = 0.28), and a random-effects meta-analysis [91] of per-locus slopes gave −0.0003 (95% CI −0.0040 to 0.0035; p = 0.90; IZ = 70%; unweighted mean −0.0002), with 9 of 20 loci negative and SORT1 contributing the largest negative slope (Figure 3b). With shuffle-adjusted BDM, full composition control and the motif and noise covariates in one model, clustered by locus, the coefficient was −0.011 (95% CI −0.022 to 0.000; cluster-robust p = 0.046; wild cluster bootstrap p = 0.11).

### 9.6 What the result does and does not establish

The pooled association survives adjustment for composition, motifs or noise covariates taken one at a time with errors clustered by dataset, shrinking by up to half, but it is not significant under inverse-variance weighting, with covariate effects allowed to differ by locus, or with the adjustments combined under locus-level inference, and it is absent on average within loci. Within loci, the falsification condition of Prediction (1), that complexity descriptors are explained by composition, GC and conservation, is therefore met in these data, and the stronger version, that complexity ranks bases within an element, is not supported either. The analysis does not show practical predictive utility, superiority to conservation, generalization beyond the 21 regulatory elements measured in reporter constructs, or any property of endogenous chromatin. Its value for the framework is that it converts the first prediction from a proposal into a tested claim that is not supported at the variant level, and it sets the scale that the genome-wide test described in Section 8.3 should expect: in these data, any pooled association is no larger in magnitude than the composition-adjusted estimates of about 0.01 to 0.02 log₂ units of effect per standard deviation of BDM, against a mean absolute effect of 0.17 log₂ units.

## 10. Limits of the Computational Frame

We list six limits of the framework.

### 10.1 Approximation looseness in algorithmic information theory

Kolmogorov complexity is uncomputable, and every empirical estimator (LZ, BDM, normalized compression distance) is a loose upper bound that depends on the choice of compressor and the alphabet [9]. The recent demonstration that the assembly index is formally equivalent to a context-free-grammar LZ compression scheme [22,23] underscores the point: measures that appear to capture fundamentally different kinds of structure may in fact be measuring the same statistical regularities by different routes.

### 10.2 Turing-completeness is a low bar

P-systems, lambda-calculus formulations of biochemistry, and several other cellular-computation formalisms are Turing-complete. This is sometimes presented as a substantive result. It is not. Almost any system with feedback and unbounded resources is Turing-complete, including systems that are biologically uninteresting.

### 10.3 Simplicity bias is contested

The claim that nature exhibits an algorithmic-probability-driven simplicity bias, phenotypes generated by short programs are more likely to evolve, is theoretically attractive and has empirical support in some genotype–phenotype maps [53]. It is also actively disputed: the bias is sensitive to the choice of universal Turing machine, the reference distribution over genotypes, and the granularity at which phenotypes are individuated [54].

### 10.4 The grammar metaphor breaks at chromatin

Formal-language descriptions of DNA assume a one-dimensional tape read in linear order. Chromatin operates on a three-dimensional contact graph, where adjacency is determined by folding rather than primary sequence. Some progress has been made on graph-based formalisms, but they are not yet at the level of expressiveness or tractability that string grammars offer.

### 10.5 Energy models are calibrated where biology is simple

Thermodynamic occupancy models work best in prokaryotes and in cell-free or reporter systems where the active components are limited and the chromatin context is absent. Mammalian enhancer logic, where most of the medically relevant non-coding variation lives, involves cooperative protein assemblies, chromatin remodeling, looping, and post-translational modifications whose parameters are not yet measured at the resolution required for predictive thermodynamics.

### 10.6 Composability across layers is not guaranteed

The argument of this paper is that the layers compose: a sequence change at layer 1 propagates through information content (layer 2), runtime context (layer 3), compiled rules (layer 4), and execution dynamics (layer 5) to produce a phenotype. Empirically this composition is partial. Sequence variants of identical algorithmic and thermodynamic profile can produce very different phenotypes depending on factors not captured by the layered abstraction, somatic mosaicism, environmental history, stochastic developmental events.

## 11. Conclusion: Testable Predictions and Falsification Conditions

The layered computational decomposition of the genome is a useful organizing frame to the extent that it makes predictions a purely correlational view does not. We close with five such predictions, each with a stated falsification condition.

1. Compressibility predicts regulatory activity independently of sequence content. Algorithmic-complexity descriptors should retain predictive power for regulatory potential when controlled for k-mer composition, GC content, and conservation. Falsification: a controlled study finds that LZ or BDM descriptors are entirely explained by these covariates. Section 9 reports a first test on measured saturation-mutagenesis data: a small pooled association does not survive full adjustment with locus-level inference and is absent within loci, so in these data the prediction is not supported at the variant level. As stated, the prediction also does not fix a sign; a stronger version should.
2. SCFG-derived structural priors improve variant pathogenicity prediction in non-coding RNA. The pipeline of Section 2.3 should improve over sequence-only baselines for clinical variant classification in ncRNA loci. Falsification: matched comparisons show no improvement after appropriate multiple-testing correction.
3. Thermodynamic occupancy models extend predictively from prokaryotic to simple eukaryotic systems given sufficient parameter measurement. Falsification: with measured binding energies and factor concentrations, occupancy models predict the measured activity of mammalian enhancers (held-out RZ or Spearman ρ on saturation-mutagenesis or MPRA data) no better than a composition-matched baseline at any cooperativity setting.
4. Rule-based reconstruction of signaling networks identifies the causal chain from a sequence variant to a phenotypic change. Falsification: for variants with a measured cellular phenotype, perturbing the intermediate nodes the model names (for example by base editing or CRISPR interference) reproduces or blocks the phenotype no more often than perturbing matched nodes the model does not implicate.
5. Small gene-regulated NCAs reproduce the developmental robustness properties of real organisms. Falsification: under matched perturbation classes (cell ablation, signaling noise, change in tissue size), NCAs with tens of genes recover the target pattern at markedly lower rates than the corresponding embryos, with recovery measured as pattern-reconstruction error relative to the unperturbed state.

Section 8.5 sets out which of these five are now directly executable against a public, genome-wide perturbation surface and which are not, and adds two further predictions, (6) grammar-class stratification and (7) the epistasis gap, that bear specifically on the boundary between precomputed variant-effect prediction and mechanistic modeling. Predictions (1) and (3) are the ones whose falsification conditions can now be attempted immediately; prediction (5) is untouched by the new resource, which is itself informative about where the execution layer still has to be built rather than looked up.

None of these predictions assumes that the genome “is” a program. They assume that treating it as one provides a useful set of formal tools whose predictions can be checked.

## Methods

### Data

Saturation-mutagenesis MPRA measurements [60] were obtained from the authors’ public release (elements.tsv.gz; https://github.com/kircherlab/MPRA_SaturationMutagenesis). The table lists every variant twice, once in GRCh37 and once in GRCh38 coordinates; GRCh38 rows were retained (44,658 of 89,316). Deletions were removed, leaving 39,170 single-nucleotide substitutions, and variants supported by fewer than ten barcodes were removed, leaving 38,556 variants across 29 element-by-condition datasets.

These represent 21 regulatory elements at 20 distinct loci: SORT1 (three datasets), TERT (four cell contexts), PKLR (two time points), LDLR (two replicates) and ZRS (two conditions) each contribute several datasets over a shared interval, and the two MYC enhancers (rs6983267 and rs11986220, about 118 kb apart at 8q24) were treated as a single locus. The response was the absolute value of the per-variant coefficient reported by the original study, a log₂ effect on the RNA/DNA ratio.

### Reference sequence and conservation

hg38 sequence for each element with 300 bp of flanking sequence was retrieved from the UCSC Genome Browser REST API [65]; the reference allele reported for every retained variant matched the retrieved base (100.00%, 0 mismatches). Per-base phyloP100way scores [63] were retrieved from the same API for every assayed position (100.00% coverage).

### Sequence descriptors

For each assayed base, windows of 21, 51 and 101 bp centered on that base were extracted. LZ76 was computed as the number of phrases in the Lempel–Ziv 1976 parse [61], divided by window length. BDM was computed with pybdm (one-dimensional, four-symbol coding-theorem-method table) [62] and divided by window length. Shannon entropy was computed over mononucleotide frequencies, and the zlib ratio as compressed length at level 9 divided by window length. GC fraction and the 64 trinucleotide frequencies were computed in the same windows; trinucleotide frequencies were standardized and reduced to their first ten principal components per window. All predictors were standardized to zero mean and unit variance before modeling.

### Statistical analysis

Models were ordinary least squares with element fixed effects and cluster-robust standard errors grouped by element (29 clusters), fitted with statsmodels 0.15. The primary model regressed absolute effect on element, GC, ten trinucleotide principal components, phyloP100way and BDM in the 51 bp window. Each of the twelve descriptor-by-window combinations was fitted with and without the conservation term, and the twelve conservation-adjusted p values were corrected together by the Benjamini–Hochberg procedure [64]. For the permutation null, BDM values were circularly shifted by a random offset within each element 1,000 times and the primary model refitted. Leave-one-locus-out analyses removed all datasets sharing a locus and recomputed principal components and standardization on the remaining data. Within-element analyses residualized both absolute effect and BDM on GC, phyloP100way and the ten principal components separately within each element, computed the Spearman correlation of the residuals, and tested the 29 correlations with a two-sided sign test and a Wilcoxon signed-rank test.

### Robustness analyses

For the robustness analyses the pipeline was re-implemented from the public data in Python 3.13 with statsmodels 0.15, pybdm 0.1 and pyjaspar 4.0. GRCh38 sequence was retrieved from the Ensembl REST API [92] and phyloP100way values from the CADD v1.7 annotation service [93], because the UCSC API could not be reached from the analysis environment; reference alleles matched at all retained variants, and the re-implementation reproduced the reported estimates to within 0.0001. Full composition control used mono-(linear and quadratic), di-and trinucleotide frequencies in the 51 bp window, reduced to orthogonal components explaining 99.5% of their variance; GC was omitted because it is a linear function of mononucleotide composition. Shuffle-adjusted BDM and LZ76 were the observed values minus their means over 50 dinucleotide-preserving shuffles of the same window [88]. Motif covariates used the 879 JASPAR 2024 CORE vertebrate profiles [89] as log-odds matrices against a uniform background, with per-profile score thresholds at p < 10⁻⁴ from the exact score distribution: the largest score loss and gain among sites matched in the reference or alternative sequence, membership in any matched site, and the number of matched profiles. Per-variant standard errors were derived from the reported coefficients and p values. Noise covariates were log barcode count and this standard error; inverse-variance weights used the standard error floored at its 1st percentile; signal-to-noise was the absolute coefficient divided by its standard error, capped at 50; and significance calls (reported p < 10⁻⁵) were modeled as a linear probability. Single adjustments kept the primary model’s clustering by dataset. Models clustered by locus, collapsed to one value per position, with locus-specific covariates, or combining shuffle-adjusted BDM, full composition control and the motif and noise covariates used errors clustered by locus; the locus-specific model interacted GC, the trinucleotide components and conservation with locus indicators. Wild cluster bootstrap p values used Rademacher weights over loci with the null imposed (9,999 draws) [90]. Per-locus slopes were estimated with locus-specific covariates, descriptors standardized within each locus and standard errors clustered by position, and combined by DerSimonian–Laird random-effects meta-analysis [91]. Code and numerical results are provided as Supplementary File 2.

### Software and reproducibility

Python 3.11 with numpy 2.4, pandas 3.0, scipy 1.17, statsmodels 0.15, scikit-learn 1.8, pybdm 0.1 and matplotlib 3.10. The four-symbol one-dimensional coding-theorem table distributed with pybdm does not contain every 12-symbol block, so BDM is an approximation; Shannon entropy, which has no such dependency, gives the same sign and comparable magnitude at every window. Scripts that reproduce every number and Figure 2 from the public data are provided as Supplementary File 1.

## Author Contributions

A.M.S. and L.P. jointly conceived the layered framework and wrote and revised the manuscript. A.M.S. performed the worked analysis in Section 9. Both authors approved the final version.

## Data and Code Availability

The saturation-mutagenesis MPRA data analyzed in Section 9 are publicly available from Kircher et al. [60] (https://github.com/kircherlab/MPRA_SaturationMutagenesis). hg38 reference sequence and phyloP100way conservation scores were retrieved from the UCSC Genome Browser REST API [65]; for the robustness analyses they were retrieved from the Ensembl REST API [92] and the CADD v1.7 annotations [93]. All code needed to reproduce the analysis and Figure 2 is provided as Supplementary File 1, and code for the robustness analyses and Figure 3 as Supplementary File 2. No other new data were generated.

AlphaGenome Atlas is available for non-commercial research use through its public web portal and the AlphaGenome API, and the AlphaGenome model is available for academic use under its own license [55,56,59]; the analyses proposed in Section 8 require no data beyond those resources and the ENCODE 4 registry [66].

## Supplementary Material

Supplementary File 1 (Supplementary_File_1_code.zip): scripts for data cleaning, sequence and conservation retrieval, descriptor computation, all statistical analyses and Figure 2, with the numerical results in JSON. Supplementary File 2 (Supplementary_File_2_robustness.zip): scripts for the robustness analyses in Section 9.5 and Figure 3, with the numerical results in JSON.

## Acknowledgements

We thank Kircher and colleagues for releasing the saturation-mutagenesis data in full.

